# Automated Classification of Sleep-Wake States and Seizures in Mice

**DOI:** 10.1101/2023.04.07.536063

**Authors:** Brandon J. Harvey, Viktor J. Olah, Lauren M. Aiani, Lucie I. Rosenberg, Danny J. Lasky, Benjamin Moxon, Nigel P. Pedersen

## Abstract

Sleep-wake states bi-directionally interact with epilepsy and seizures, but the mechanisms are unknown. A barrier to comprehensive characterization and the study of mechanisms has been the difficulty of annotating large chronic recording datasets. To overcome this barrier, we sought to develop an automated method of classifying sleep-wake states, seizures, and the post-ictal state in mice ranging from controls to mice with severe epilepsy with accompanying background EEG abnormalities. We utilized a large dataset of recordings, including EMG, EEG, and hippocampal local field potentials, from control and intra-amygdala kainic acid-treated mice. We found that an existing sleep-wake classifier performed poorly, even after retraining. A support vector machine, relying on typically used scoring parameters, also performed below our benchmark. We then trained and evaluated several multi-layer neural network architectures and found that a bidirectional long short-term memory-based model performed best. This ‘Sleep-Wake and Ictal State Classifier’ (SWISC) showed high agreement between ground-truth and classifier scores for all sleep and seizure states in an unseen and unlearned epileptic dataset (average agreement 96.41% ± SD 3.80%), and saline animals (97.77% ± 1.40%). Channel elimination and feature selection provided interpretability and demonstrated that SWISC was primarily dependent on hippocampal signals, yet still maintained good performance (∼90% agreement) with EEG alone, thereby expanding the classifier’s applicability to other epilepsy datasets. SWISC enables the efficient combined scoring of sleep-wake and seizure states in mouse models of epilepsy and healthy controls, facilitating comprehensive and mechanistic studies of sleep-wake and biological rhythms in epilepsy.

## 1. Introduction

While the relationship between sleep and seizures has been widely appreciated for centuries (Crespel et al., 1998; Janz, 1962; Shouse & Sterman, 1982), mechanisms remain obscure. As a prelude to rodent studies examining this relationship, we sought a means to comprehensively label (score) sleep, wake, and seizure-related activity in large datasets from continuous chronic mouse electroencephalographic (EEG) recordings.

Several factors hinder the meticulous investigation of sleep in rodent models of epilepsy: Firstly, there are variable changes in the EEG background and prominent interictal EEG abnormalities (Pitkänen et al., 2017). Secondly, these abnormalities are not static, typically evolving from seizure induction throughout the recording period. Lastly, there is substantial variability in the number, severity (Almeida Silva et al., 2016), and electrophysiological morphology (Henshall et al., 2000) of seizures and interictal findings, necessitating larger cohorts than might be needed for studies of sleep-wake alone. These large datasets often involve thousands of hours of multi-channel recording. Manual scoring of this data for both seizures and sleep-wake is often impractical; a trained expert scorer may take more than 25 minutes to score sleep-wake for 12 hours of data (Kloefkorn et al., 2020), and even longer when EEGs are abnormal.

Sleep-wake and seizure classification have independently been achieved. Earlier approaches to sleep classification, using linear discrimination, depend on highly simplified features such as assessing the EEG power ratio in the theta and delta bands and mean and standard deviation of EMG (Costa-Miserachs et al., 2003). Overall, these existing methods likely depend on typical EEG background rhythms and features that are disrupted in epilepsy (Kilias et al., 2018; Song et al., 2024). More complex methodologies for sleep scoring include approaches utilizing Support Vector Machines (SVMs) (Lampert et al., 2015) and machine learning-based approaches. The latter type of algorithms include several classifiers driven by convolutional feature extraction. AccuSleep (Barger et al., 2019) uses a linear classification layer, and MC- SleepNet (Yamabe et al., 2019) uses a Bidirectional Long Short-Term Memory Layer (BiLSTM) with a Dense layer for classification. Previously, techniques including convolutional feature extraction were used to score sleep and cataplexy in cataplexic mice (Exarchos et al., 2020). However, these existing sleep classifiers are trained on non-epileptic mice with a normal EEG background, and do not account for the seizures and the post-ictal state.

**S**everal effective seizure detection approaches have been published. Methods include parametric (Tieng et al., 2017), machine-learning (Wei et al., 2020), and deep-learning-based (Jang & Cho, 2019) algorithms. These previous works are also adequate for identifying seizures across various mouse models of epilepsy (Wei et al., 2021). However, like the existing published sleep scoring classifiers, none combine sleep-wake and seizure classification.

Large datasets with prolonged recording are needed to study the important phenomenological and mechanistic relationships between sleep-wake and biological rhythms. These datasets necessitate an automated way to perform combined sleep-wake and epilepsy-related classification. We aimed to create an automated sleep-wake and seizure scoring method that could batch-process and accurately score files from control mice and mice with varying degrees of epilepsy-related EEG background abnormalities. We also sought to use our sleep-wake and seizure state data to evaluate and directly compare which signal features and approaches resulted in the most accurate sleep and seizure identification. We focused on machine learning methods that are either theoretically appropriate or empirically suited to classifying EEG time series data. As a benchmark, we sought to achieve a classification accuracy comparable to that seen with human scoring, as determined by inter-rater agreement. The agreement between scorers in our laboratory is high at >93% (Kloefkorn et al., 2020), in accordance with other reports of 92% (Rytkönen et al., 2011). Here, we describe a highly accurate method for simultaneous automated sleep-wake and seizure classification, the Sleep-Wake and Ictal State Classifier (SWISC).

## 2. Methods and Materials

### 2.1. Mice

Mice (9-31 weeks old, n=79) of either sex (n=34 male, n=45 female) were used in accordance with the Emory Institutional Animal Care and Use Committee. Mice in this dataset were obtained from Jackson Labs and were wild-type C57BL/6J (n=16, n=8 of each sex, Stock No. 000664), or VGAT-ires-Cre Knock-In C57BL/6J (VGAT-Cre, n=63; n=37 female and n=26 male, Stock No. 028862). VGAT-Cre mice were used to obtain baseline sleep-wake and epilepsy data as a prelude to future studies with this genotype. The intra-amygdala kainic acid model was used for C57BL/6J mice given a relatively lower mortality rate compared to other chemical kindling approaches in this strain (Conte et al., 2020), as well as its utility for later electrical kindling experiments (Straub et al., 2020). All animals were bred in our animal facility with the oversight of the Department of Animal Resources. Breeding procedures included backcrossing every five generations. DNA samples were obtained via ear punch before surgery to determine the genotype using polymerase chain reaction per the Jackson Labs protocol. Mice were provided with food and water *ad libitum* and maintained on a 12-hour light-dark cycle (lights on 7 am-7 pm). Cages were changed weekly and in the same session for all mice. A total of 21 animals were excluded from analysis given epilepsy-related death (n=12, VGAT-Cre) or technical failure ((n=9; 6 VGAT-Cre, 3 wild-type) before the study end point three weeks after kindling.

### 2.2. Surgery

Surgical procedures have been described previously (Zhu et al., 2020). Briefly, mice were induced with ketamine (100 mg/kg, i.p.) and xylazine (10 mg/kg, i.p.), followed by meloxicam (5 mg/kg s.c.) in 1 cc saline, and anesthesia was maintained with isoflurane (0-1.5%). Four holes were drilled for head plate screws, two for depth electrodes, and one for a guide cannula targeting the basolateral amygdala. Bilateral hippocampal depth electrodes were placed in the perforant path region, immediately overlying the dorsal blade of the dentate gyrus (±2.00 mm ML, -2.53 mm AP, -1.80 mm DV from the brain surface). Using a headplate-mounted recording montage developed by our laboratory, screw electrodes for electrocorticography (ECoG) were placed in the left frontal (-1.30 mm ML, +1.00 mm AP) and right parietal bones (+2.80 mm ML, -1.50 mm AP), as well as a reference over the midline cerebellum (0 mm ML, -6.00 mm AP) and a ground in the right frontal bone (+1.30 mm ML, +1.00 mm AP). Finally, the guide cannula (5 mm, Plastics1, Roanoke, VA) was implanted with the tip 1.75 mm dorsal to the basolateral amygdala target (-2.75 or -3.25 mm ML, -0.94 mm AP, -3.80 mm DV from the brain surface). Electromyogram (EMG) paddles (Plastics1, Roanoke, VA) were inserted subcutaneously above the posterior neck muscles, and instrumentation was then secured with cyanoacrylate adhesive and dental cement. The mice were then allowed time to recover from anesthesia and regain their righting reflex before being singly housed in seven-inch diameter clear acrylic recording barrels with food and water available *ad libitum,* nesting material, and a 12-hour light-dark cycle.

### 2.3. Mouse Recording

Mice recovered from surgery for three to four days and were then connected to the tether for two days of habituation before seven days of baseline recording. Video, EEG (ECoG and bilateral hippocampal field potential [HPC-L/R]), and EMG were recorded continuously throughout the experiment. A one-megapixel day-night camera was used for video recording (ELP, Shenzhen, China). Pre-amplifying headsets (8406-SE31 M, 100x gain, Pinnacle Technologies, KS) were used with a commutator (model 8408, Pinnacle Technologies, KS) and analog breakout boxes (8443-PWR, Pinnacle Technologies, KS) with additional gain and digitization (Power 1401, CED, Cambridge, UK). Synchronized video and EEG/EMG files were saved every 12 hours and automatically restarted with Spike2 (v9.10, CED, Cambridge, UK).

### 2.4. Kainic Acid Injection

After continuous baseline recording, either kainic acid (KA; n=51) or normal saline vehicle (n=7) was injected into the basolateral amygdala 5-7 hours after lights on (0.3 μg in 200 nL over 5 minutes) via the internal cannula after removal of the stylet. Video-EEG recordings were continued throughout the KA infusion. Mice were injected with diazepam (5 mg/kg, i.p.) to terminate status epilepticus 40 minutes after the end of the KA infusion. The recording then continued for an additional three weeks.

### 2.5. Manual Sleep and Seizure Scoring

Scoring was performed by one of three authors (LR, LA, NP), and then confirmed by those with the most experience (LA, NP). Sleep scoring was performed manually by a conventional approach with visual assessment of the four recording electrographic channels, a spectrogram of EEG and hippocampal depth electrode, root-mean-squared EEG, with video recording available for disambiguation (not used for scoring, but available for Racine staging).

Each 20-second epoch was assigned one of five possible labels (Wake, rapid eye movement (REM), non-REM (NREM), seizure, or the post-ictal state, see Figure 1) when half or more than half of the epoch consisted of that state except in the case of seizure epochs. Sleep-Wake states were labeled by conventional criteria. Wakefulness is characterized by variable and predominantly theta through gamma EEG activity, often with phasic EMG activity. NREM is characterized by high delta power and loss of beta and gamma activity in the EEG, along with lower EMG activity than wakefulness. NREM sleep was not divided into sub-states as it is in humans, which is typical for rodent scoring (Rayan et al., 2022). REM is associated with lower EMG activity than non-REM, low delta EEG activity, and high theta activity, particularly in hippocampal electrodes, with the latter becoming less prominent in some epileptic mice.

**Figure 1.**
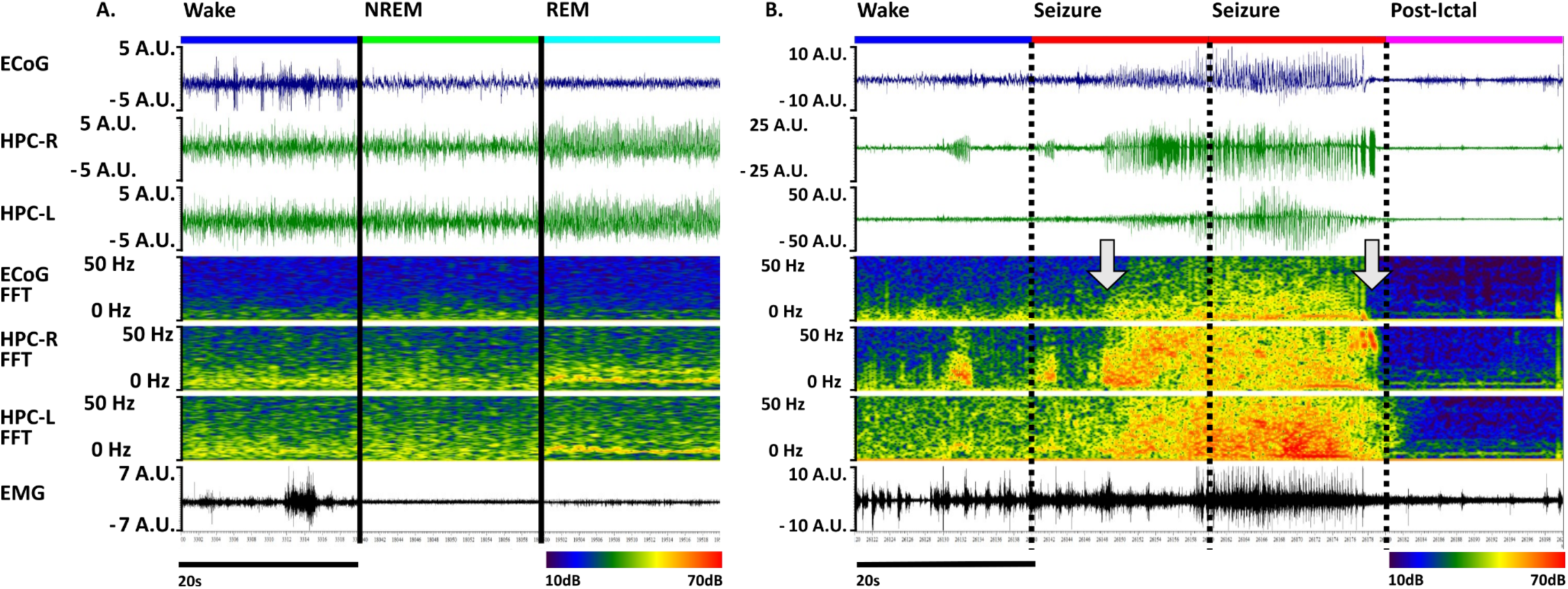
Scoring of mouse recordings. A. Sleep-wake categorization on three non-consecutive twenty-second epochs drawn from electrocorticography (ECoG), left and right hippocampus (HPC-L and HPC-R), and electromyogram (EMG) channels. Y-axes are presented in arbitrary units (a.u.) to reflect potential voltage range differences between mouse cohorts. The use of normalization in the pre-processing pipeline addresses concerns about using specified units on inputs. In the same channels, a spontaneous seizure occurs from wake in the intra-amygdala kainic acid (IAKA) mouse model, divided into twenty-second consecutive epochs, followed by post-ictal obtundation. A spike train begins in the left hippocampus during the epoch labeled “wake,” progresses into a full seizure before the midpoint of the second epoch (left arrow), and ends just before the third epoch ends (right arrow), resulting in a label of “seizure” for the second and third epochs. The fourth epoch shows the suppressed electrographic signal characteristic of a post-ictal state. The magnified image of this seizure can be found as Figure 1-1.

Scoring was adjusted based upon rules for mouse sleep adapted from American Academy of Sleep Medicine (AASM) and Rechtschaffen and Kales criteria (Moser et al., 2009; Rechtschaffen & Kales, 1968)): REM can only follow NREM, and NREM can only be scored when there are two or more consecutive epochs.

Seizure scoring was performed via visual EEG scoring that required five seconds or more of rhythmic spikes that were evolving in morphology, frequency, or amplitude (see Figures 1 and Supplementary Figure 1). Epochs that contained the bulk of the seizure, for short seizures, or included more than five seconds of seizures were scored as ‘seizure’. The rationale for this is that seizures are typically of high spectral power and dominate the epoch’s normalized feature vector (see below). Seizures in rodents typically dominate the EEG signal, making sleep-wake scoring fraught, and are likely associated in many cases with impaired awareness. Thus, seizures were scored as a distinct state from sleep-wake states. The post-ictal state is characterized by initial behavioral obtundation and post-ictal electrographic suppression, but can be scored without reference to video. This state is marked and remits suddenly, so visual scoring was used (see below for agreement between scorers). This state was included given that it is a state of behavioral obtundation and/or forebrain dysfunction that does not fit into typical sleep-wake states.

### 2.6. Dataset Composition

The dataset, including unscored data, contains 3,770 files (1,885 days) from experimental KA mice and 650 files (325 days) from either pre-treatment baseline recordings or saline-injected controls. Of those, a total of 900 files (450 days) from experimental KA mice and 279 files (139.5 days) from either pre-treatment baseline recordings or saline-injected controls had been manually scored (∼27% of the dataset). Data containing either uninterpretable recording errors (i.e., values that are “not a number” [NaNs] after computation) or text-encoded markers containing sleep scores that were not part of the target scoring states (such as markers for scoring on which experts disagreed) were not used to ensure error-free computation. Three files were excluded by these rules, leaving a final total of 1176 files.

### 2.7. Computational Resources

Machine learning was implemented using the Python libraries TensorFlow (Google, version 2.10.0; Abadi et al., 2016) and Keras (version 2.10.1; Chollet, 2018), with GPU training and inference using cuDNN version 8.1 and Cudatoolkit version 11.2.2 (NVIDIA et al., 2021). These versions were used to ensure forward Windows compatibility for future development of end-user tools. The class distribution (Data Distribution, section 2.10) contained several imbalanced classes, which were accounted for using the SciKitLearn (version 1.0.2; Pedregosa et al., 2011) compute_class_weight function (2.13. Statistics and Classification Metrics). All file conversion, feature extraction, model training, and model inference were performed on a desktop PC (Intel i7-6950X at 3.00 GHz, 128 GB RAM, Nvidia RTX4090 24GB).

### 2.8. Preprocessing

A custom script exported files for each mouse from Spike2 into MATLAB format. A Python Jupyter notebook was then used to perform the following preprocessing steps. Files were imported from the .mat format using HDF5Storage (version 0.1.18), then data was filtered using a first-order Butterworth high-pass filter at 1 Hz, using Scipy (version 1.7.3). Z-scoring was then performed to normalize amplitudes on a per-file basis. Decimation was performed using Scipy’s signal package to resample the data from 2 kHz to 200 Hz, including a second-order infinite impulse response anti-aliasing zero-phase filter for ECoG, and an eighth-order version of the same filter for the HPC-L, HPC-R, and EMG channels. Numpy (version 1.21.6) was used to order the data into an array of 20-second epochs and save the data in the .npy format. Feature extraction was then performed via Numpy’s real Fast Fourier Transform (Frigo, 1999)) and the Scipy Signal Welch Power Spectral Density function (see 2.9. Feature Extraction below and Figure 2).

**Figure 2.**
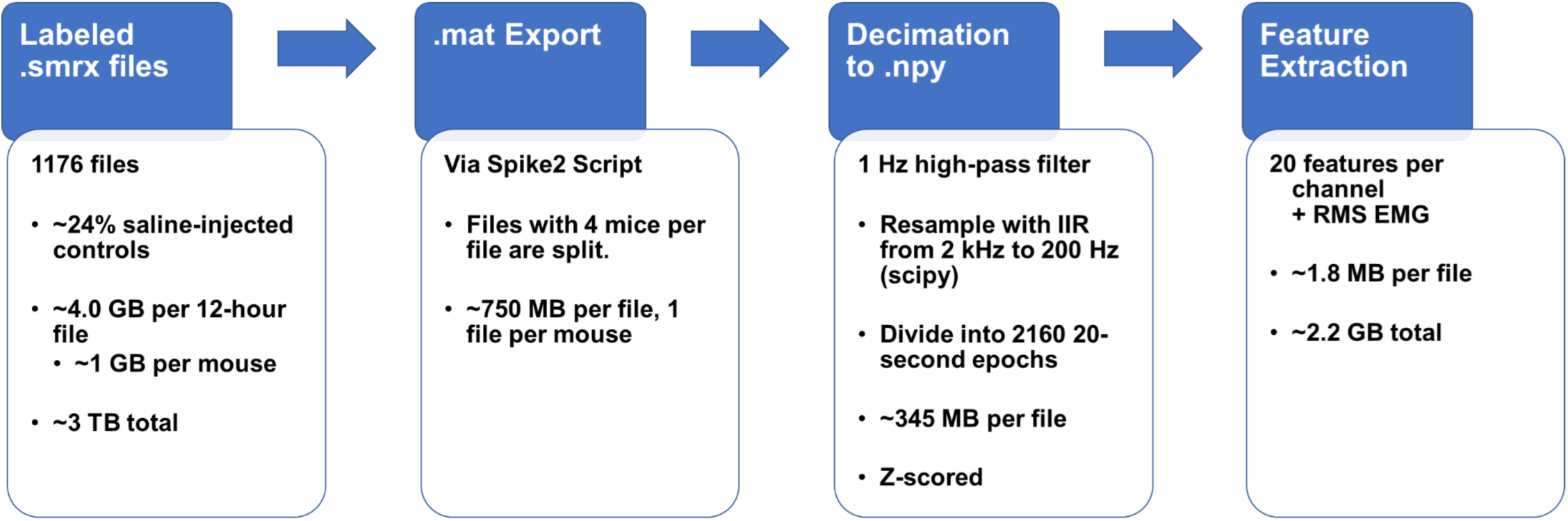
Data Pre-Preprocessing Pipeline in Python. Recorded data, as in Figure 1, is stored as .smrx, an EEG-specific file format. These files are large and not easily read into Python, they must first be exported with 20-second epochs to MATLAB .mat format via a script built in Spike2, the recording and analysis software. The .mat files are then downsampled by a factor of 10, including a zero-phase infinite impulse response anti-aliasing filter implemented in Scipy, then z-scored for normalization and exported to .npy to further save disk space. Finally, feature extraction is performed, described further in Feature Extraction.

### 2.9. Feature Extraction

Six statistical features for each channel were extracted regarding the time-domain information of the signal within each epoch: mean, median, standard deviation, variance, skewness, and kurtosis. These statistical features were selected for their physiological relevance: Median, standard deviation, and variance are commonly extracted features for EEG analysis (Stancin et al., 2021). Skewness and kurtosis were chosen as additional features due to their specific relevance and demonstrated effectiveness in EEG signal processing in epilepsy (Xiang et al., 2020).

Additionally, spectral features were calculated for each canonical frequency range of the EEG: delta (δ, 2-4 Hz), low theta (θ, 4-7 Hz), high θ (7-13 Hz), beta (β, 13-30 Hz), low gamma (γ, 30-55 Hz), and high γ (65-100 Hz). While line noise was low, given the recording configuration, we excluded 55-65 Hz to ensure this classifier did not have line noise-related problems if used in other settings. We computed the absolute magnitudes of the real portion of Fast Fourier Transforms (FFT) and Welch Power Spectral Density (PSD). Both FFT and PSD were used, as PSD is normalized to the width of the frequency range over which it is calculated. This provides a more accurate gauge of the relative power in bins of differing frequency widths, as we have in our paradigm. Each of these power estimates was normalized to the broadband FFT or PSD in the 2-55 Hz range to ensure that all spectral power measures were normalized relative to the baseline power of the epoch of interest. A ratio of delta power to low theta power was also calculated for both FFT magnitude and PSD, paralleling a primary feature used for manual scoring (often called “theta-delta ratio” or theta:delta). In total, 20 features were gathered for each of the four channels. Finally, 20 one-second root-mean-squared (RMS) amplitude values are calculated from EMG for each epoch and concatenated with the other channel features for a total of 100 features per epoch.

The above features were then split into groups which were compared to validate our feature selection and provide some direct comparison to manual scoring where applicable. The first consisted solely of the four channels’ theta:delta in both FFT magnitude and PSD and the full vector of RMS EMG amplitude for the epoch. This closely reproduces the commonly used features for manual sleep-wake scoring. This first feature set we refer to as “Delta/Theta and RMS” (DT/RMS). As previous classifiers based primarily on these features have worked in non-epileptic animals, this was a key feature to include to demonstrate the need for a more advanced classifier for the analysis of kainic-acid-treated animals. Feature Set 2 consisted solely of the 6 statistical features for each channel in the epoch, and RMS EMG - a total of 44 features. Feature Set 3 included only the four channels’ Fourier magnitudes in the selected bins, the delta/theta ratio, and the RMS EMG. No epoch-level statistical features nor Power Spectral Density were used for Feature Set 3, referred to as FFT/RMS. The final evaluated feature set contained all statistical features, FFT and PSD magnitudes, delta/theta ratios, and RMS EMG components of the full 100-feature vector and is referred to in subsequent text and figures as the Full feature set. See Figure 2-1 for a visual summary of tested feature sets and frequency bins of interest.

Extracted features for each epoch were input to ScikitLearn and Keras models in array format, with each feature vector in the input *x* array corresponding to an epoch label at the same index in the *y* array. As epochs were exclusively scored as one of five states, *y* label arrays were encoded as one-hot labels with 5 indices. These indices correspond to [wake, NREM, REM, seizure, post-ictal]. For example, an epoch scored as wake would be represented as [1,0,0,0,0].

### 2.10. Training, Validation, and Test Dataset Creation

The data were fully separated into training/validation/testing datasets on a per-subject basis to avoid any in-sample training. Data used for training contained animals of all types, with ∼97.3% being VGAT-Cre KA animals (n=34) and ∼2.7% being VGAT-Cre saline control animals (n=1). The overrepresentation of VGAT-Cre KA animals was intentional, as this phenotype has marked inter-subject variability in the power of frequency bands typically used to detect sleep. The validation dataset used to assess loss functions during training contained solely VGAT-Cre saline control animals (n=3), to assess the generalizability of the factors learned from the training dataset to saline-treated animals.The holdout testing dataset consisted of ∼35% VGAT-Cre KA (n=7), 50% wild-type KA (n=10), and ∼15% wild-type saline controls (n=3). In building this additional testing dataset from these recent cohorts, we can ensure that the classifier generalizes fully to a different genotype and to the introduction of novel cohorts of animals. Of the 1176 scored 12-hour recordings, the final file split was ∼60.12% (n=707 files) for training, 10.97% (n=129) for validation, and 28.91% (n=340) for testing the final versions of the evaluated models. This data split was effective for training with manually-scored data while reliably producing an accurate classification on completely out-of-sample validation animals.

### 2.11. Model Architectures

We selected several approaches to machine learning that were based in either what we took to be the implicit processes involved in the human scoring, mirrored non-machine learning approaches, or adopted architectures that had previously been found to be effective. We started by examining the performance of a highly effective classifier of sleep-wake in non-epileptic mice, ‘AccuSleep’, that is open-source and thus configurable for our dataset. Next, a support vector machine approach was used as a benchmark, given that a discriminant function is used or implicit in manual and spreadsheet approaches’ reliance on theta:delta and RMS EMG We hypothesized this would not perform well for epileptic mice, despite reasonable classification for controls given disrupted theta:delta due to background slowing and disrupted theta in epileptic mice. The other approaches were based upon multi-layer neural network models that had previously been shown to be effective for automated sleep scoring in humans or rodents. To compare methods, we trained several varieties of these models on our three sets of extracted features, then compared the performance of these models with validation and test datasets to determine the best model for our application.

#### 2.11.1. AccuSleep

We first wanted to determine how an effective sleep-wake classifier would perform with epileptic mice. We used the open-source AccuSleep framework (Barger et al., 2019), given that it could be directly compared to the structure of our data and re-trained as necessary. Briefly, AccuSleep is a multilayer convolutional neural network that classifies images of the spectrogram of log-normalized spectra for each epoch, in addition to normalized RMS EMG. Labels of wake, NREM, REM, or unknown are assigned with customizable epoch lengths. AccuSleep can be downloaded pre-trained with various epoch lengths and used with custom epoch lengths by pooling shorter epochs (we pooled 10 s epochs to create 20 s epochs). We tested this pre-trained model with the 34 manually scored files that included seizures from the testing dataset. Files with seizures were chosen to exclusively evaluate the classifier’s performance on recordings with epilepsy-induced electrophysiological abnormalities. We then retrained AccuSleep with 20 second epochs and manually-scored labels from all files containing seizures from our training dataset (109 files), and then compared to pre-trained performance with the 34 seizure-containing files from the testing dataset.

#### 2.11.2. Support Vector Machine

The baseline architecture for comparison to human scoring was a Support Vector Machine (SVM) architecture (Vapnik, 1997) with a linear discrimination function. Support Vector Machines, simply, take a set of (*y, x)* points, *y* is a class label and *x* is a feature vector, and map them to points (*y, z)* in a higher-dimensional feature space *Z*, where the label *y* corresponds to a derived set of features *z* in the high-dimensional space. The classification problem is then solved by the support vector machine, by determination of a hyperplane which maximally separates the sets of points within this higher-dimensional feature space. The SVM used herein relies on a hinge loss function for multiclass classification (Crammer & Singer, 2001). The SVM, trained with an iteration count of 1000 using SciKitLearn’s linear_model.SGDClassifier function with L2 regularization was trained and validated against all four feature sets.

#### 2.11.3. Multi-Layer Architectures

We selected four multi-layer architectures for classification that were either commonly used architectures, or had been shown as effective for sleep-wake classification: (1) Dense (Fully Connected) Layers: Dense layers are the simplest starting point for a neural network, being fully-connected layers that return the dot product of their received inputs and the weights learned by the layer’s kernel. Dense layers used herein operated on a linear activation function. The use of this architecture is a baseline, as this is a widely-used and relatively straightforward architecture, and its success or failure in classification is used to evaluate the necessity for more complex architectures. Dense Layer architectures were implemented with a Dense layer (of variable size) to perform the tensor operations on the input sequences, a Flatten layer used to compress the sequences from 3 dimensions to 2 dimensions, and a 5-way softmax output layer used in the grid search for hyperparameter tuning. Long Short Term Memory (LSTM): LSTMs are a type of Recurrent Neural Network (RNN) where three gates (input, output, and forget), along with input from previous time steps, are leveraged to control which learned weights are remembered from past predictions and used to predict the current time step (Hochreiter & Schmidhuber, 1997). LSTM Implementation: LSTM architectures were implemented with an LSTM layer (of variable size), a 40% Dropout Layer for regularization, a Flatten layer, and a 5-way softmax output layer. Bidirectional Long Short Term Memory (BiLSTM): BiLSTMs are a variant of LSTM layers that perform their forget-gate operations on time steps in both the forward and backward directions, as opposed to the solely backward-looking operation of LSTMs. This provides much more utility in the context of predicting labels in cases where the signal characteristics of time points later in the signal are known, as is the case in offline vigilance and ictal state scoring (Graves & Schmidhuber, 2005). BiLSTM architectures were implemented with a BiLSTM layer (of variable size), a 40% Dropout Layer, a Flatten layer, and a 5-way softmax output layer. We also implemented a stacked variant of the BiLSTM (Stacked BiLSTM), with each of four BiLSTM layers halving in size as they progress. The first layer in the chain is of variable size. The final BiLSTM layer was then used as input to a 40% Dropout layer, a Flatten layer, and a 5-way softmax output layer.

### 2.12. Grid Search Paradigm for Multi-Layer Architectures

Training was executed using each possible combination of various manually specified parameters to tune the model’s inputs and features. The first feature tested in our grid search of models was the three feature vectors. The second feature tested in this grid search was the number of units in the variable-size base layer of each architecture at variants of 50, 100, and 200 units. For testing purposes, the same layer size input was used in LSTM as in BiLSTM, resulting in the first layer of the BiLSTM architectures consisting of [2 x layer size] input units, where layer size is denoted in figures and tables. Our third feature evaluated was input sequence length, where the epoch of interest was input in sequence with several preceding and following epochs. The assessed variants of input sequence length were 1, 3, 5, and 7. For example, at sequence length 1, epoch *xT* is paired with epoch state label *yT.* At sequence length 7, epoch xT is presented to the classifier as the vector {xT-3, xT-2, xT-1, xT, xT+1, xT+2, xT+3}, with the state label *yT*. The use of input sequences of varying lengths is standard with the use of LSTM classification and is the equivalent of a sliding-window feature extraction approach.

The initial grid search consisted of a combinatorial search of four feature vectors, four machine learning models, three-layer sizes, and four input sequence lengths. With the addition of the four feature vectors tested for the SVM evaluation, the total number of models evaluated in the initial grid search was 196. To reduce computational time, 20 training epochs using the full training data were performed for the first round of evaluation before classification matrices and reports were saved. The early stopping criterion for all training was decided to be when there was a lack of change of .001 in loss function in 5 training epochs. Examples of layer architectures, with all assessed variables represented in the architecture diagrams, are presented in Figure 3 and the accompanying legend.

**Figure 3.**
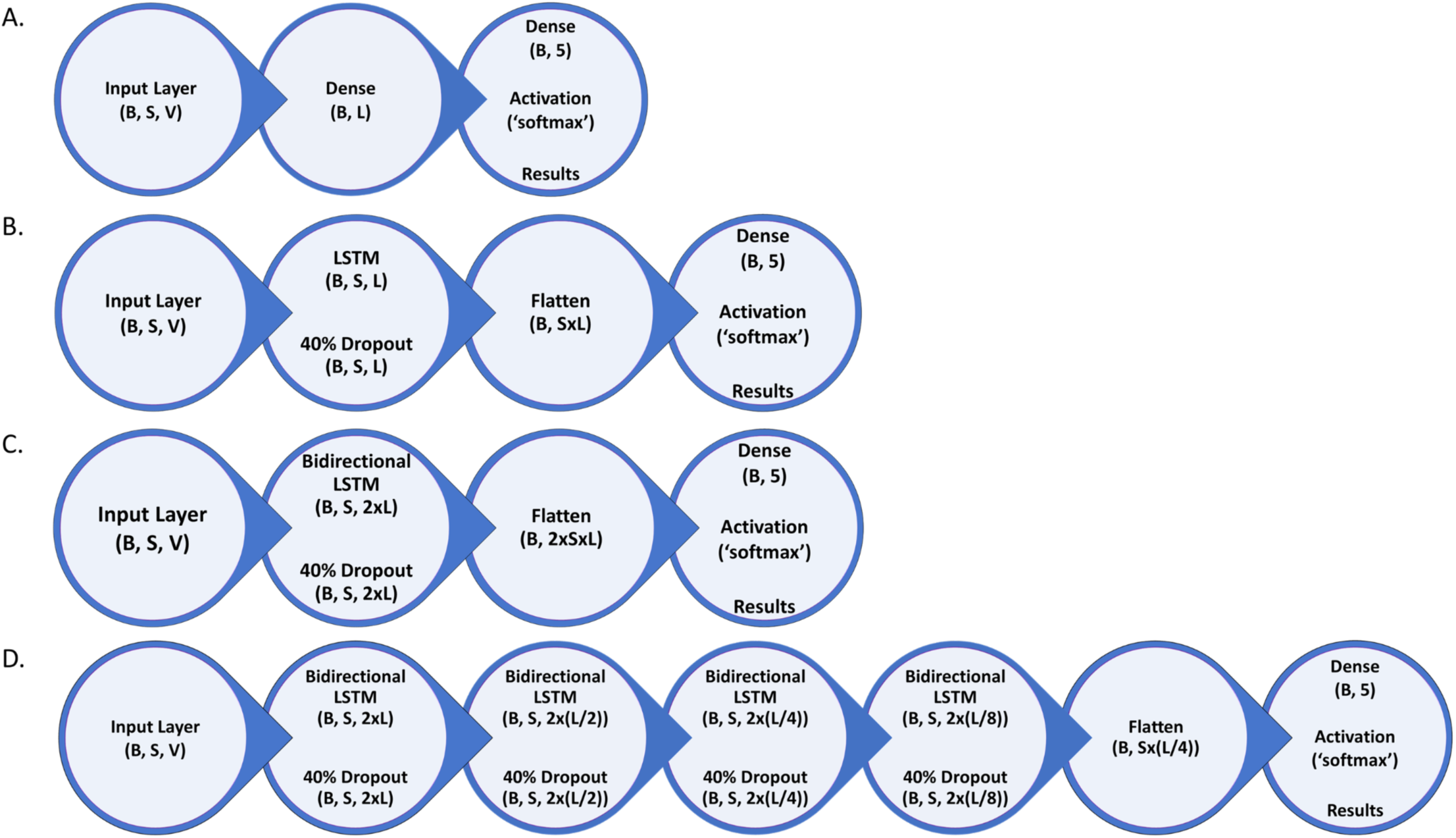
Layer Architecture Search Space. Graphical representation of the layer architectures tested and the parameters applied. Variables tested in the grid search are represented here with single letters. B represents the batch size of 2160, that is, the length in 20-second epochs of one full 12-hour electrographic recording. S represents the variable sequence length of epochs in the input vector. V represents the variable length of the input vector itself.. L represents the variable layer sizes of 50, 100, or 200.

### 2.13. Statistics and Classification Metrics

A modified version of the output from the SciKitLearn compute_class_weight method was used to provide class weight inputs for the imbalanced classes to Keras. The weights generated by this function are used as input to the class-weight parameter of Keras, and are used to assign a relative weight to each class with respect to their impact on the loss function of the machine learning model. The compute_class_weight function was used with the parameter “balanced,” via the following equation:

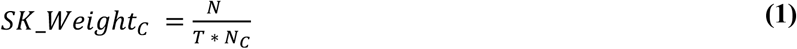

In equation 1, N denotes the number of samples, NC denotes the number of samples of a given class, T denotes the total number of classes, and SK_WeightC is the value returned for that class by the compute_class_weight method. This class weighting function, however, gives a very large range of values for classes where the counts differ by orders of magnitude, causing Keras to operate inefficiently. To overcome this limitation, each number in the resulting class weight array was then modified via the following equation to smooth the values in the class weighting array while maintaining their relative scales.

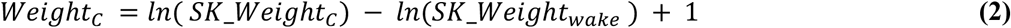

Equation 2 sets the weight of wake to 1, and scales the rest of the weights relative to ln(SK_Weightwake).

Using Keras’s built-in metrics methods the following classification metrics were calculated for each epoch of training: true/false positives, true/false negatives, categorical accuracy, precision, recall, area under the precision-recall curve (AUCPR), and categorical cross-entropy loss.

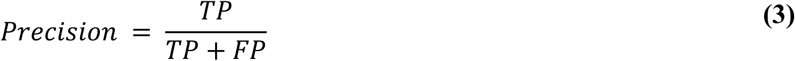

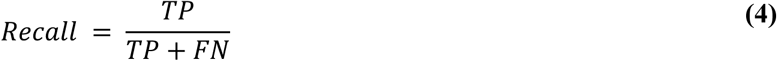

The AUCPR is approximated as the Riemann sum of a plot of precision versus recall values at 200 different thresholds, all of which are calculated for each one-hot encoded state for a given epoch. Categorical cross-entropy loss is a loss function used to optimize and evaluate the classifier. This loss function is calculated by the cross-entropy function (Liu et al., 2020).

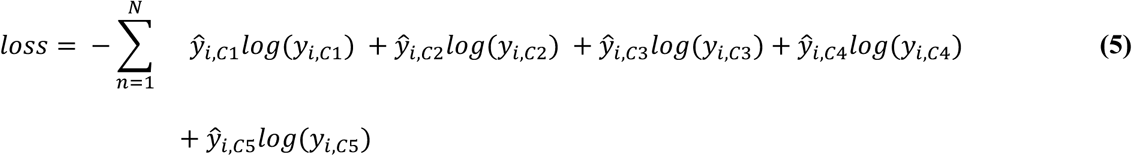

In equation 5, *yi,C1* to *yi,C5* represent each of the labels in the one-hot encoding for a given sample, and *ŷi,C1* to *ŷi,C5* represent the five outputs from the 5-way softmax output layer. In all of the machine learning models used, we optimize the loss function using the Nadam optimizer (Dozat, 2016).

SciKitLearn’s built-in metrics methods were also used to produce true/false positives, true/false negatives, categorical accuracy, precision, and recall for the purposes of output to Excel format. SciKitLearn also allowed for the calculation of the F1 score for each class given by the following equation:

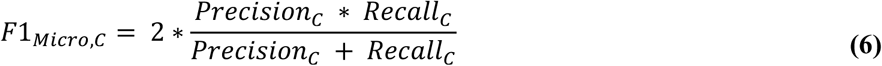

In equation 6, *F1Micro.C*, *PrecisionC*, and *RecallC* denote the score calculated for that class. These scores were calculated via the SciKitLearn classification_report function using the true and predicted labels for each epoch, determined by the class with the highest-scoring prediction probability. F1Micro scores will be used for cross-model analyses except in cases where precision or recall are substantially different from, and thus not properly summarized by, F1Micro.

The macro and weighted multiclass F1 scores were also calculated by the SciKitLearn classification_report function according to the following equations, where NClasses is the total number of classes identified, and N is the total number of samples:

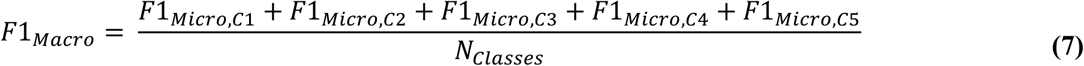

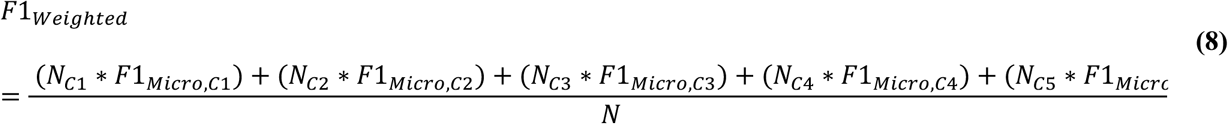

Confusion matrices were created using SciKitLearn’s confusion_matrix function.

### 2.14. Classification Performance of Trained Classifier with Shorter Epochs

After optimal model selection based on the grid search, we sought to examine generalization of this classifier to shorter epochs. We selected four-second epochs. Four second epochs are the common lower bound of rodent sleep scoring. These short epochs are the lower bound of feasible manual scoring and still allow spectral estimates for low frequency EEG activity. Twenty second epochs are often used when raw amounts of sleep-wake are studied; four second epochs are more appropriate when examining sleep fragmentation or narcolepsy models with brief cataplexy. The change of epoch length required a modification to the signal preprocessing to adapt RMS EMG to the feature vector. The RMS EMG was calculated as before, in one-second steps. This vector of four one-second bins, {RMS1, RMS2, RMS3, RMS4}, was then distributed to the 20-feature RMS EMG vector with 5 repeats per RMS value. This repetition allows the four second signal to fit the existing 20-second feature vector.

## 3. Results

### 3.1 Performance of Existing Sleep-Wake Classifier (AccuSleep) in Epileptic Mice

We hypothesized that existing sleep-wake classifiers would underperform with epileptic mice given interictal EEG abnormalities including with epileptiform discharges and alterations in the EEG background. We evaluated a highly effective classifier that could be modified to work with our data, and retrained with data from epileptic mice. We used the pre-trained classifier with modified epoch lengths (see Methods) to classify 20-second epochs as wake, NREM, REM, and unknown. *Precision* scores for the pre-trained AccuSleep were: wake: 0.966; NREM: 0.414; REM: 0.297. *Recall* scores for the pre-trained AccuSleep were: wake: 0.356; NREM: 0.471; REM: 0.736. *F1Micro* scores for the pre-trained AccuSleep were: wake: 0.520; NREM: 0.440; REM: 0.423. While positive classifications (precision) were high for wake, identification of all epochs (recall) was low, and the overall performance was not acceptable for sleep-wake scoring (Figure 4A and C).

**Figure 4.**
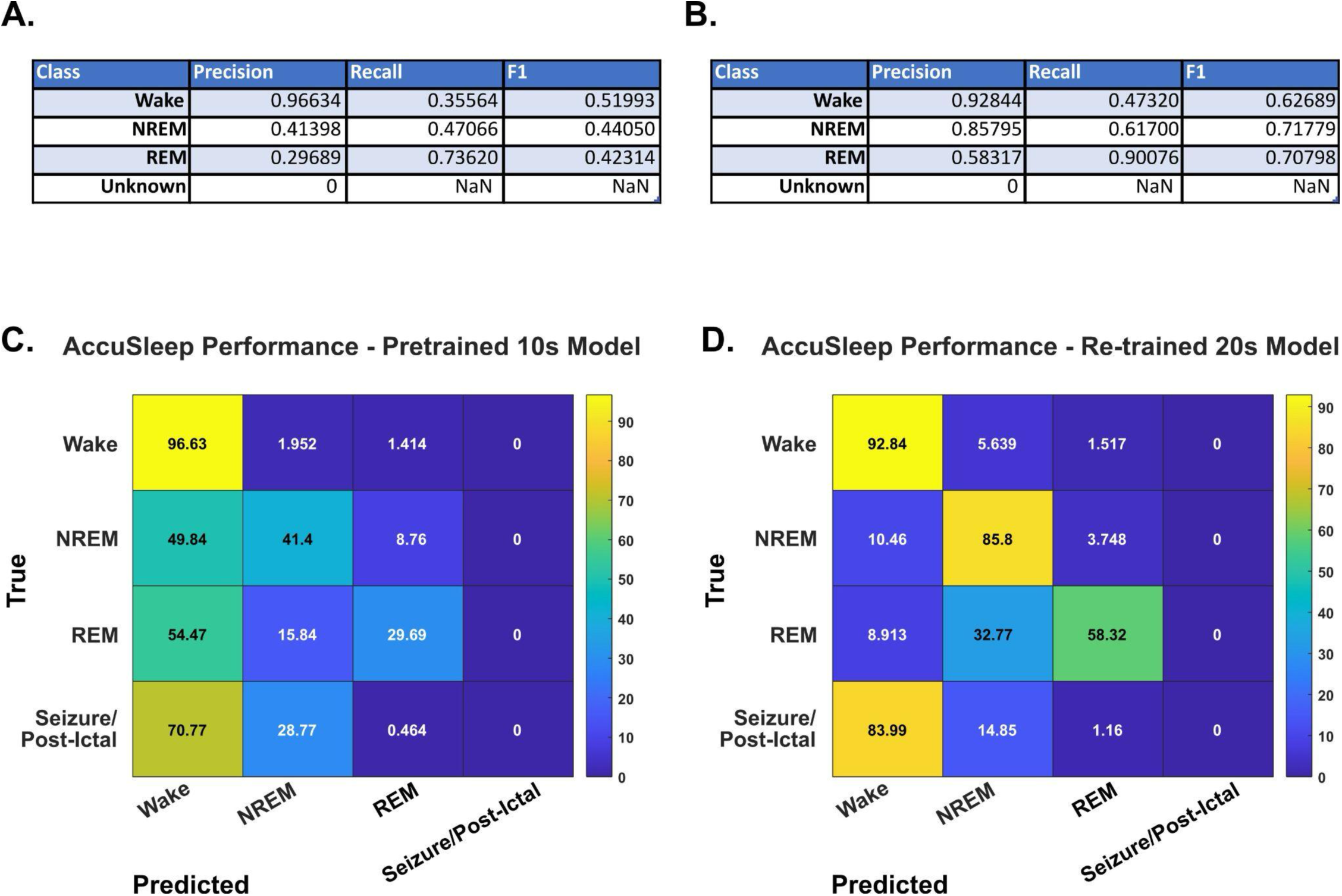
AccuSleep Performance on Epileptic Mouse Data. A.-B. Precision, Recall, and F1 values for each state for the 10-second-epoch (A.) and 20-second epoch (B.) AccuSleep classifier. This classifier does not have the ability to train or score seizure states, so these were given as “Unknown” for the ground truth. B. The confusion matrix of a pre-trained 10-second-epoch AccuSleep classifier on 34 recording files featuring all sleep-wake states as well as seizures from the cohorts used in the testing dataset described in Methods. It is evident that, while the classifier can precisely classify wake states, it does not have proper recall and is prone to false-positive wake states for both NREM and REM classes. The confusion matrix of a customized version of AccuSleep trained on 109 recording files to evaluate 20-second-epochs. These recording files were the files from the cohorts used in the training dataset described in Methods, which featured sleep-wake states as well as seizures. This version of the classifier can precisely identify wake states as well as NREM states, but is prone to false-positive REM states for NREM epochs. This classifier also identifies 14% of seizures and post-ictal states as NREM.

Using our training dataset and the provided training scripts from AccuSleep we obtained substantially better performance. *Precision* scores for the newly-trained AccuSleep were: wake: 0.928; NREM: 0.857; REM: 0.583. *Recall* scores for the newly-trained AccuSleep were: wake: 0.473; NREM: 0.617; REM: 0.717. *F1Micro* scores for the newly-trained AccuSleep were: wake: 0.583; NREM: 0.901; REM: 0.707 (see Figure 4B and D). Despite the improvement with retraining, classification of all instances of each state (recall) was still too low for our purposes and did not reach our benchmark, and particularly with regard to seizures.

For both tested versions of AccuSleep, no seizures were correctly classified with the appropriate “Unknown” label, even when a version of the classifier was specifically trained to score existing known seizure epochs as Unknown (see Figure 4). This performance of AccuSleep on our data corresponds with our suppositions about how the epilepsy associated changes in spectral character would cause existing classifiers to underperform on such animals. Given that this otherwise effective classifier underperformed for epileptic mice, we then compared some reasonable alternative approaches.

### 3.2. Support Vector Machine

Our first architecture to train on our data to determine the proper feature set was the SVM, which is most comparable to manual scoring approaches. This testing found that the DT/RMS feature set, using an SVM, produced inferior classification precision, and a near-zero recall and *F1Micro* score for REM [*Precision:* 0.175*, Recall:* 0.008, *F1Micro*: 0.0170] as compared to both wake [*Precision:* 0.646*, Recall:* 0.862, *F1Micro*: 0.738] and NREM [*Precision:* 0.641*, Recall:* 0.862, *F1Micro*: 0.738] in the mixed-treatment testing dataset. In addition, seizure state precision, recall, and *F1Micro* scores were zero, and all post-ictal classification measures were zero. The Stat/RMS feature set improved Wake [*Precision:* 0.822, *Recall*: 0.882, *F1Micro*: 0.851], NREM [*Precision:* 0.784, *Recall*: 0.773, *F1Micro*: 0.778] and REM classification [*Precision:* 0.651, *Recall*: 0.241, *F1Micro*: 0.352], however seizure and post-ictal both had unacceptable seizure [*Precision:* 0.572] and post-ictal classification [*Precision*: 0.183]. The FFT/RMS feature set improved classification for the testing dataset in all states: Wake [*Precision:* 0.936*, Recall:* 0.936, *F1Micro*: 0.936], NREM [*Precision:* 0.897*, Recall:* 0.911, *F1Micro*: 0.904], and REM [*Precision:* 0.750*, Recall:* 0.672, *F1Micro*: 0.709], seizure [*Precision:* 0.516, *Recall*: 0.276, *F1Micro*: 0.360] and post- ictal [*Precision:* 0.318, *Recall*: 0.401, *F1Micro*: 0.355]. Finally, the Full feature set, when evaluated against the testing dataset, performed well for wake [*F1Micro*: 0.935], and NREM [F1Micro: 0.900], with lesser results for REM [*F1Micro*: 0.744], seizure [*F1Micro*: 0.825], and post-ictal [*F1Micro*: 0.426] classes. Overall, the SVM performed well for some states, but is still below benchmark, even with improved performance with the full feature set (see Figure 5).

**Figure 5.**
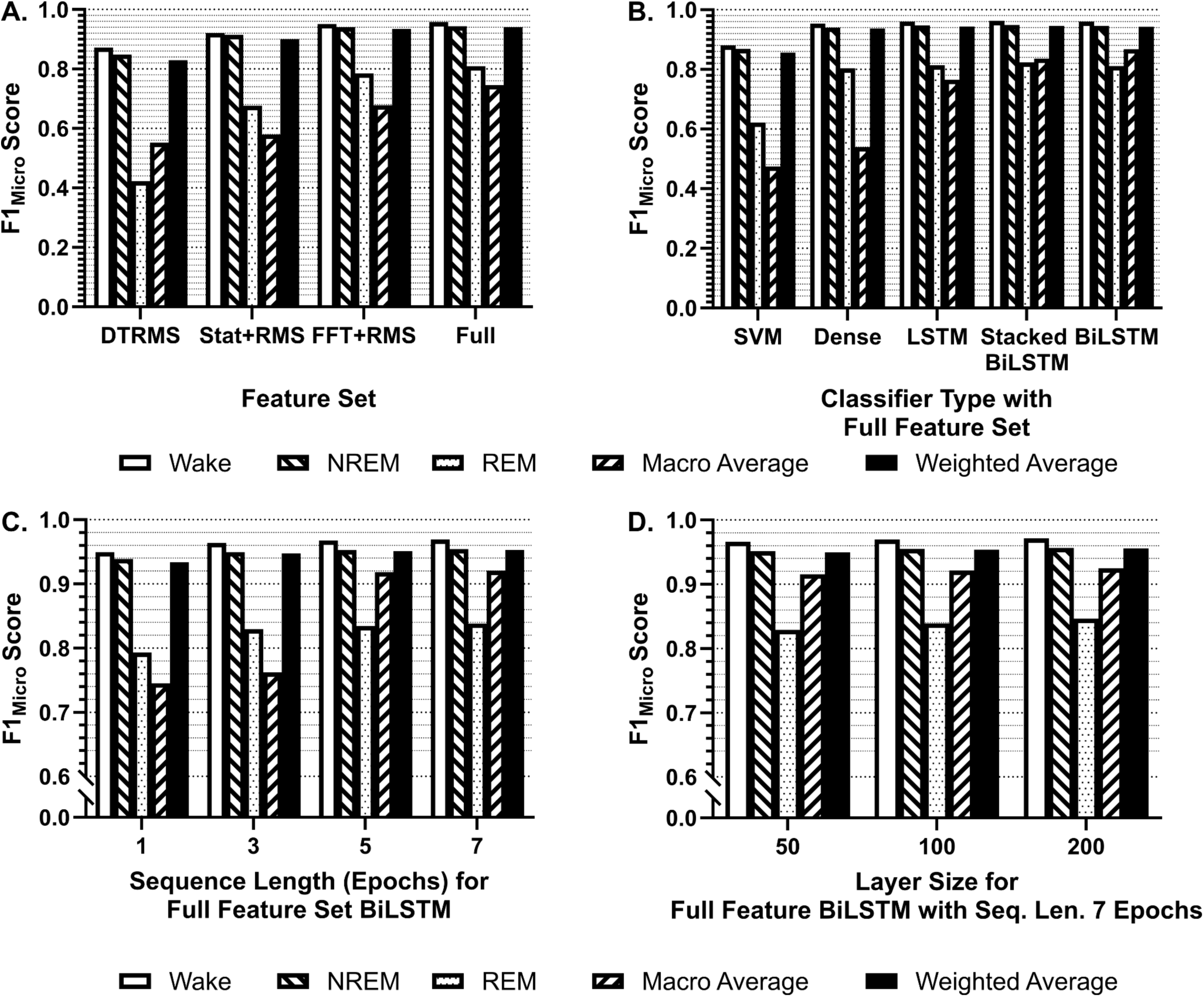
F1 Metrics at 20 Epochs of Training. *F1Micro* scores for the individual states contained in the validation dataset (wake, NREM, REM) as well as *F1Macro* and *F1Weighted* were assessed at each nested level of the grid search to determine the best-performing model. Each feature set, architecture, sequence length, and base layer unit count were exhaustively validated against one another. A. For each feature set tested, each of the *F1* metrics is averaged over all architectures, sequence lengths, and layer sizes. At this level of analysis, the greatest performance across all metrics was the Full feature set. B. For each architecture tested using the Full feature set, each of the *F1* metrics is averaged over sequence length and layer size. At this level of analysis, the greatest performing classifiers were of the BiLSTM architecture. LSTM and Stacked BiLSTM were comparable in performance. C. For each sequence length tested as input to a BiLSTM architecture using the Full feature set, each of the *F1* metrics is averaged over all layer sizes. At this level of analysis, the greatest performing classifiers were trained with a sequence length of 7. D. For each base layer unit count tested using 7 sequence length inputs to a BiLSTM architecture using the Full feature set, each of the *F1* metrics is averaged over all layer sizes. At this final level of analysis, the best-performing classifier on the test dataset in the 192-model grid search is the 200 base unit count, 7 sequence length input, BiLSTM classifier using the Full feature vector.

### 3.3. Multi-Layer Architectures

We next compared four multi-layer network architectures, as described above: Dense layer, LSTM, BiLSTM, and Stacked-BiLSTM (see Figure 3). We compared these with 20 training epochs (passes through the training dataset, not to be confused with data epochs). The distribution of classification metrics varied substantially by class. Focusing on *F1Micro, F1Macro*, and *F1Weighted* as well-rounded metrics described previously, we evaluated the *F1* scores for all classifiers across our classes to determine the best-performing classifiers tested in our grid search. We evaluated the impact of feature vectors, classifier architecture, sequence length, and base layer unit size. After training, our results from ranking all parameters determined that the Full feature vector, BiLSTM architecture, 7-epoch input sequence, and 200-unit base layer size were the best-performing parameters in each respective category. This architecture, stopped at 20 epochs of initial training, achieved F1Micro scores of 0.972 on wake, 0.957 on NREM, and 0.846 on REM, with an *F1Macro* of 0.925 and *F1Weighted* of 0.956 on the saline validation dataset. On the holdout testing dataset, this classifier achieved an F1Micro on wake of 0.978, on NREM of 0.958, on REM of 0.887, on seizure of 0.782, and post-ictal of 0.741, with an *F1Macro* of 0.869 and *F1Weighted* of 0.965 (see Figure 5).

While the BiLSTM performance was best, we next investigated somewhat similar performance of LSTM, BiLSTM, and the Stacked-BiLSTM architectures with increased training. All three classifiers were retrained with the 7-epoch input sequence, a 200-unit base layer size, and the Full feature vector from the beginning, this time with a training limit of 60 training epochs or to an early stopping threshold of 0.001, as defined in the Methods. *F1Micro, F1Macro*, and *F1Weighted* metrics achieved by these models on the holdout testing dataset were used for the final evaluation. *F1Micro* scores for the training, validation, and testing sets across all states, as well as *F1Weighted* for these models are ranked and presented in Figure 6.

**Figure 6.**
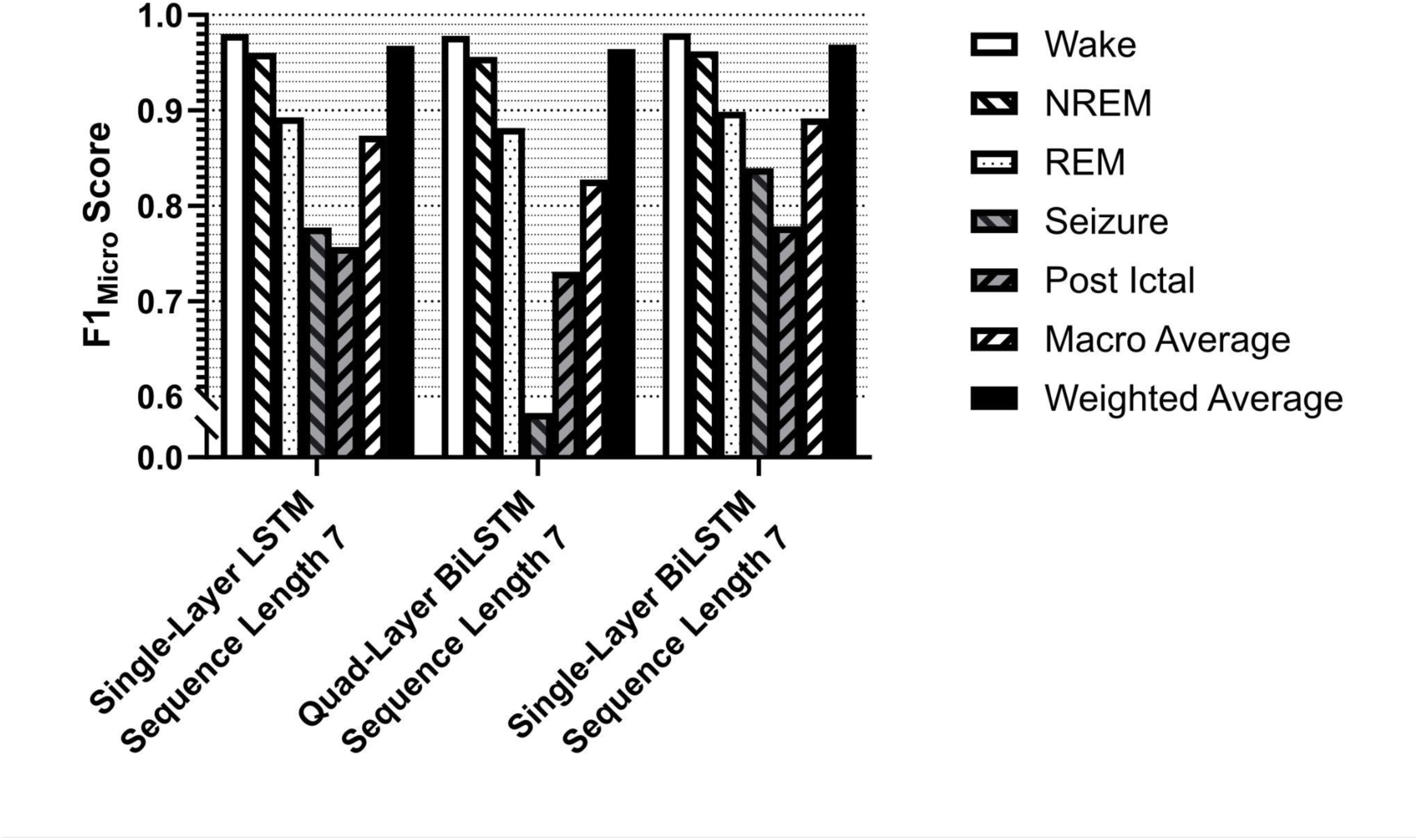
F1 Metrics After Complete Training. *F1Micro* scores for the individual states contained in the unseen and unlearned testing dataset (wake, NREM, REM, seizure, post-ictal) as well as *F1Macro* and *F1Weighted* were assessed for all of the LSTM-based variants trained with the optimized parameters: the Full 100-feature vector, an input sequence length of 7, and 200 base layer units. These three resulting architectures were trained to a limit of 60 epochs, with a loss patience of .001 over 5 epochs. Ultimately, the classifier that performed the best against the holdout real-world testing dataset was the Single-Layer BiLSTM, achieving an *F1Weighted* of 0.968, an *F1Macro* of 0.886, and an *F1Micro* of the seizure state of 0.824.

Our final evaluation of these three classifiers found that a classifier with a 7-sequence-length input vector using the Full feature space with an architecture consisting of a 200-unit base layer single BiLSTM with a 40% Dropout and a 5-way softmax output layer was the most effective for classification of sleep as well as seizure and post-ictal state (Figure 7, denoted as 7-Full-BiLSTM). This comprised our final product, the Sleep-Wake and Ictal State Classifier (SWISC). *F1Micro* scores for the SWISC for our validation dataset were 0.974 for wake, 0.959 for NREM, 0.860 for REM, with an *F1Macro* of 0.931 and an F1Weighted of 0.959. In the case of the holdout testing dataset, *F1Micro* scores for the SWISC were 0.981 for wake, 0.962 for NREM, 0.898 for REM, 0.840 for seizure, and 0.778 for post-ictal, with an *F1Macro* of 0.891 and an *F1Weighted* of 0.968 The final weighted accuracy across classes for this model was 96.59%. Confusion matrices produced for all training/validation/testing sets and all classes for the fully-trained SWISC are presented in Figure 8, showing the true classification of wake, NREM, REM, and seizure at or above 90%, with variation in the post-ictal state accounted for its more qualitatively defined nature described earlier in this manuscript. The complete breakdowns of all metrics for all models tested can be found on *github.com/redacted*.

**Figure 7.**
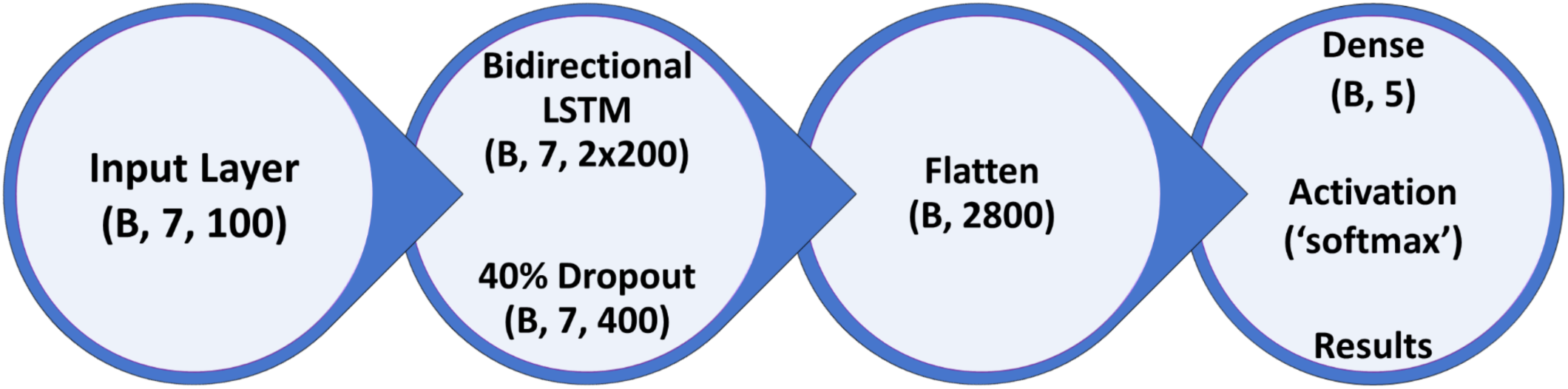
Final Model Architecture. The classification section of the model consists of an input layer for epoch sequences, followed by a BiLSTM layer of 200 units in each direction. The generalization of the classifier is improved by using an activity regularization function with an L1 of .0001 on the BiLSTM layer, then a 40% Dropout layer. The output of this section is then flattened, and classification is performed by a 5-unit Dense layer with softmax activation.

**Figure 8.**
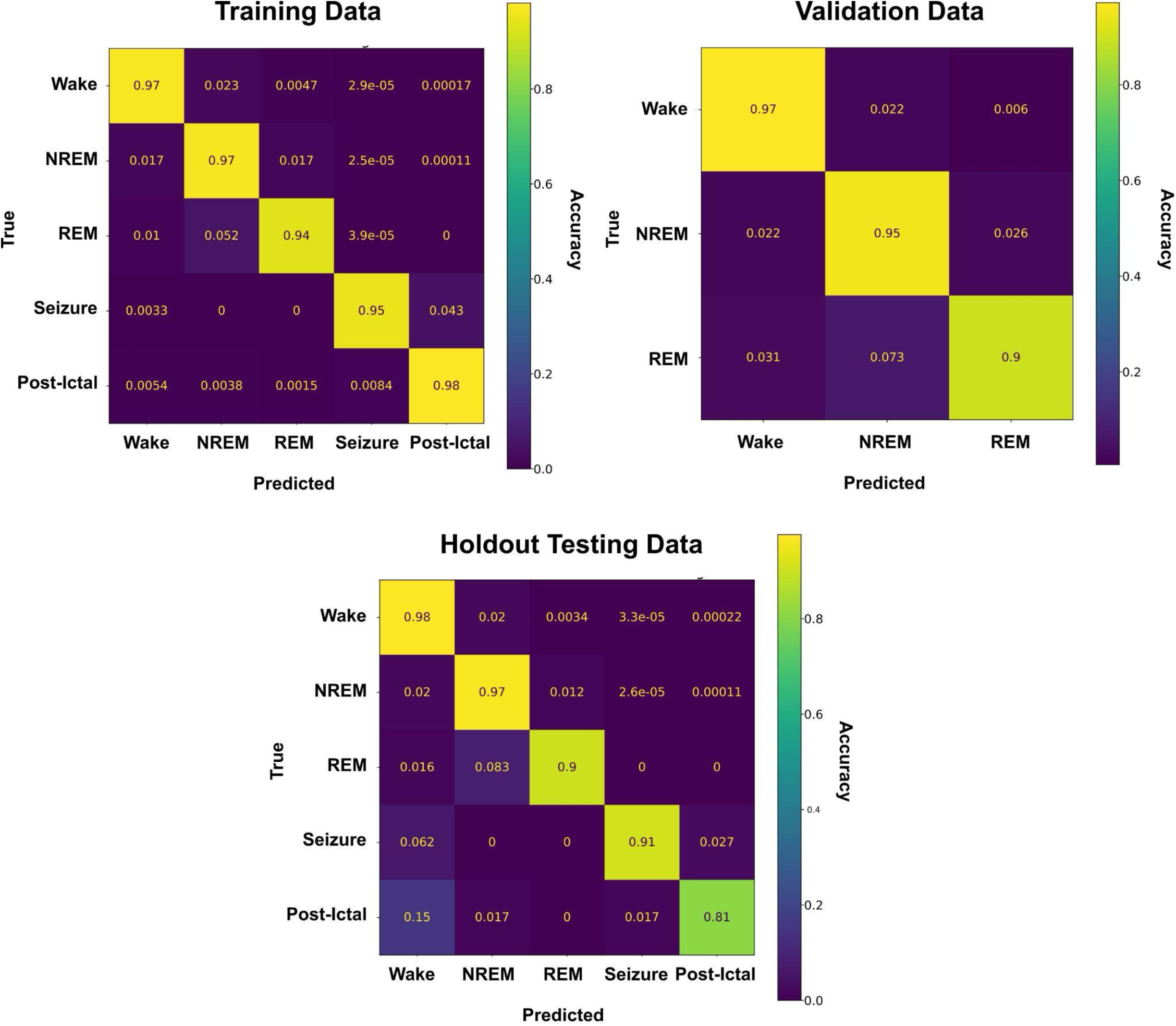
Performance after 60 Training Epochs. With the 2x200-unit initial layer size, the training and validation datasets were scored accurately compared to expert scoring (low 90%), mirroring its performance on the training and control datasets without overfitting.

### 3.4. Interpretability - Channel Dropping

To better understand the individual electrophysiological channels’ contribution to the chosen architecture’s scoring accuracy, we systematically removed individual channels. Then we trained new instances of the SWISC model out to 60 epochs or early stopping to properly evaluate these performances against the testing dataset.

Scoring based on hippocampal channels alone was highly effective. Training this architecture with solely our bilateral hippocampal channels, using all 20 statistical and spectral features present for each channel, produced *F1Micro* scores on our testing dataset for wake at 0.964, NREM at 0.952, REM at 0.866, seizure at 0.818, and post-ictal at 0.872, with an F1Macro of 0.914 and F1Weighted of 0.949. This shows that even with half of the original classifier’s inputs, robust classification is possible using this architecture. Adding EMG statistical and spectral features and RMS EMG to the dual-hippocampal montage produced similar *F1Micro* scores, with wake at 0.975, NREM at 0.959, REM at 0.888, seizure at 0.833, and post- ictal at 0.882. F1Macro and F1Weighted in this condition were 0.923 and 0.962, respectively. It is also notable that when trained on the left hippocampal channel alone (the side of kainate injection), the classifier achieves F1Micro results in wake of 0.962, NREM of 0.952, REM of 0.817, seizure of 0.841, and post-ictal of 0.813. *F1Macro* and *F1Weighted* in this condition were 0.860 and 0.947.

Scoring based on the input of the ECoG channel alone retained useful classification. *F1Micro* scores in the testing dataset of this variant still reached usable levels with wake at 0.966, NREM at 0.941, REM at 0.757, seizure at 0.829, and post-ictal at 0.779. *F1Macro* and *F1Weighted* here were 0.840 and 0.945, respectively. The saline validation dataset in this variant achieved sleep classification *F1Micro* scores of wake at 0.905, NREM at 0.931, and REM at 0.918, showing that the expansion of this classifier to sleep studies in much simpler montages in non-epileptic mice is achievable, broadening the potential reach of this classifier even further.

The addition of EMG spectral and RMS features to the ECoG-only feature vector did not improve classification, in fact reducing the F1Micro of seizure to 0.786 and post-ictal to 0.542 for the testing dataset.

As expected, scoring on EMG spectral and RMS features alone poorly discriminated between forebrain-related states. This version of the classifier still achieved high *F1Micro* scores for wake of 0.946 and NREM of 0.856, but faltered for REM with an *F1Micro* of 0.484. Seizure had an *F1Micro* of 0.271, and the *F1Micro* of the post-ictal state was 0.308. While EMG could crudely classify sleep and wake states, it displayed poor performance with REM, seizure, and the post-ictal state. Confusion matrices for the testing dataset for all of these interpretable machine-learning variants are displayed in Figure 9.

**Figure 9.**
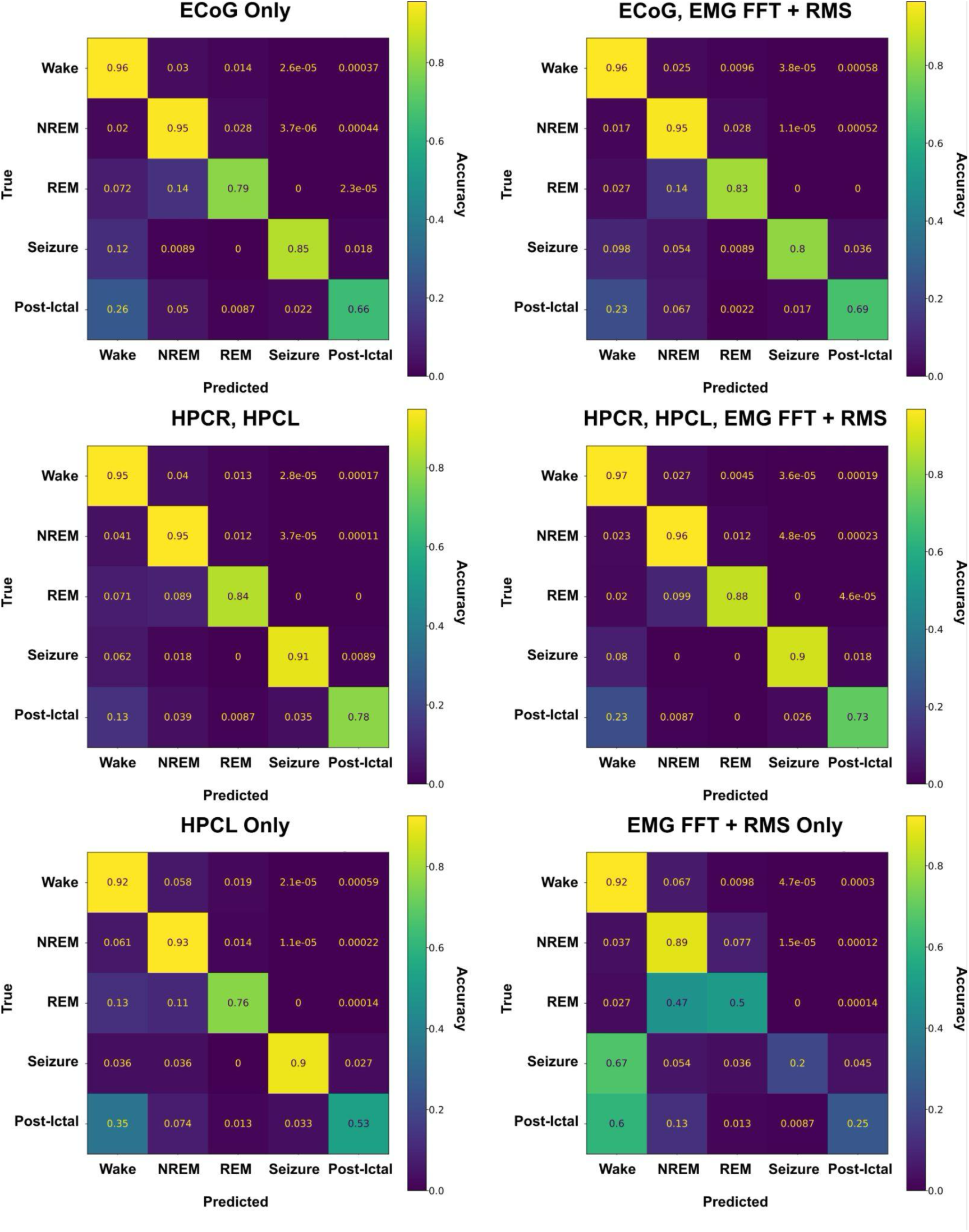
Interpretable Machine Learning via Masking. Masking specific data channels during training, a hands-on method of interpretable machine learning, showed reliable classification with the winning model when used in any configuration that included a hippocampal channel. Classification accuracy did not drop significantly unless classification was performed without hippocampal signal or ECoG. This demonstrates that the classifier shows promise for various recording montages.

### 3.5 Scoring Results Comparison

When ground-truth scores from manually scored components of the testing dataset are compared to those produced by the SWISC and rated for agreement across time epochs, average classification agreement with our ground-truth expert scores for epileptic mice in the holdout testing dataset is 96.41% (standard deviation ± 3.80%) when all states are accounted for. Holdout saline animals show an average agreement of 97.77% (standard deviation ± 1.40%). The average agreement across our full dataset of epileptic mice is 96.76% (standard deviation ± 3.30%). Additionally, there is a 96.38% (standard deviation ± 3.91%) agreement between ground-truth and classifier scores across recordings from saline-treated mice regardless of dataset. Figure 10 contains a graphical representation of agreement and the corresponding hypnogram from expert scores and the classifier for an individual file to support the classifier’s accuracy visually.

**Figure 10.**
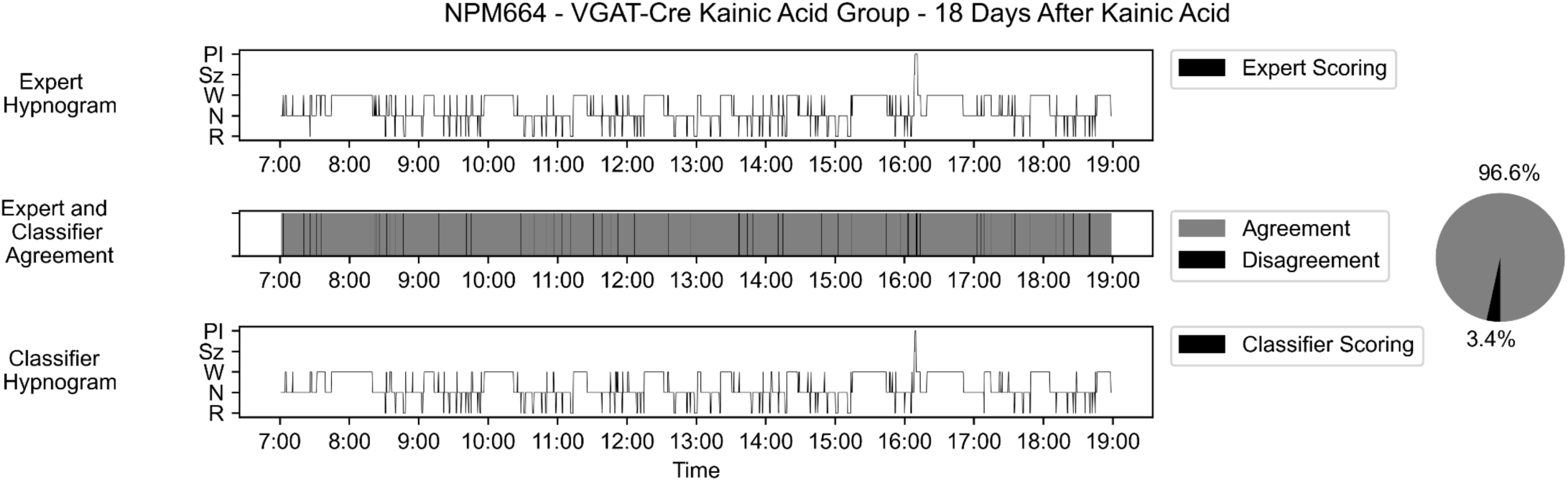
Scoring Comparisons. To test the classifier, we chose to visually and computationally compare the agreement between expert scorers and the classifier for a representative 12-hour record of sleep-wake. In a KA-treated animal, the classifier performs with ∼96% overall accuracy relative to expert scoring, which is comparable to inter-rater reliability on sleep-wake scoring tasks as demonstrated in Kloefkhorn et al., 2020, where 93% agreement was reported between 3 expert scorers. Hypnogram legend for states: PI: post-ictal; Sz: seizure; W: wake; N: NREM; R: REM.

Scoring results from the SWISC were also evaluated according to Rechtshaffen and Kales’ criteria regarding state transitions (Section 2.5). The evaluation of our automatic scoring for these two rules found only 0.20% of the total epochs scored consisted of violations of these rules, with 26.55% of these violations being lone epochs of NREM and 73.44% being REM transition violations. This suggests that the classifier may have learned to implement some form of the Rechtshaffen and Kales rules. The Rechtschaffen and Kales layer is therefore provided, as it was in another paper from this lab (Exarchos et al., 2020) to correct any such violations.

### 3.6. Performance on Shorter Epochs

A key final test for ensuring that our classifier is generalizable to other sleep analysis paradigms is ensuring that it works with differing epoch lengths. To this end, the files from our dataset were pre-processed with an epoch length of four seconds, as described in the pre-processing and feature extraction subsections. These differences in pre-processing and extraction were performed to test the limits of the classifier’s accuracy. Average agreement for 4-second epochs was assessed via manual scoring of 10 selected files, with this selection including two files per animal, one 3 days before and one 6 days after the day of kainic acid or saline injection. Of these animals, one was from the saline group, and four were from the kainic acid group. All files were scored by both BH and either NPP or DL. Inter-rater agreement for 4-second manual scores was 97.01% (standard deviation ±1.04%). The agreement of these scores with the 4-second classifier scoring was 93.81% (standard deviation ±3.24%) for BH and 93.74% (standard deviation ±3.00%) for NP/DL. The four-second scoring also largely respects the Rechtschaffen and Kales criteria, with a violation rate of 0.64%, with 50.1% being lone NREM epochs and 49.9% of these being wake-REM transitions.

## 4. Discussion

We present the successful creation of a machine-learning-based classifier for the automated, accurate, and rapid scoring of sleep-wake states, seizures, and post-ictal states in mice with epilepsy. While previously reported classifiers effectively classified sleep in various populations of mice (Exarchos et al., 2020; Grieger et al., 2021; Lampert et al., 2015; Yamabe et al., 2019), none have demonstrated scoring proficiency in any rodent models of epilepsy. We also showed that even a highly effective sleep classifier (AccuSleep) does not perform well even when re-trained to classify sleep-wake in training data from epileptic mice. As demonstrated by our grid search and comparison to AccuSleep, our BiLSTM architecture also empirically outperforms the CNN of AccuSleep, as well as SVMs, LSTMs, and Dense Neural Net classifiers trained on the exact same data.

To our knowledge, our classifier is the first to achieve the goal of combined sleep-wake and seizure classification in mice with a variable phenotype from control to severe epilepsy. This thereby overcomes the infeasibility of comprehensive sleep-wake classification in epileptic mice that limited study of the important bidirectional interactions of sleep and epilepsy (Bernard et al., 2023; Sheybani et al., 2025). This classifier may have broad applicability given that classification performance remains high without EMG (common in studies of epilepsy), and even with ECog or hippocampal LFP alone. However, we were not in a position to test the classifier with other epilepsy models that may have markedly different electrophysiological features, such as absence models, and markedly different electrophysiologic phenotypes seems likely to require re-training of the classifier.

With our classifier, scoring time can be reduced to less than 3 minutes per 12-hour recording file, including all pre- processing steps. There is no human input or time needed other than visual and statistical assessment of the results. These time savings will make the analysis of larger datasets feasible, as is often required in epilepsy studies due to the large individual variations in epilepsy severity. The time savings are also not traded off for any loss of precision in the sleep scores. The scoring accuracies of the classifier, with a weighted average of >95%, exceed the 93% agreement in human scores (Kloefkorn et al., 2020) for AASM/Rechtshaffen and Kales sleep scoring between scorers in our lab, making it a well-rounded classifier for sleep and equivalent to a trained human scorer. Although seizures in rodents are variably operationalized, inter-rater agreements are high in our experience, as is the performance of our classifier.

The network architecture chosen fits well with the design intent. The innovation of BiLSTM is that it adds further information to the classifier by adding prior and future epochs in time series analysis rather than classifying the epoch at hand without sequence information (Graves & Schmidhuber, 2005). This likely accounts for the implicit inclusion of Rechtshaffen and Kales scoring rules behavior. On the other hand, the need for past and future epochs limits the use of the classified for immediate closed-loop control.

In summary, this classifier provides a rapid, accurate, robust, multi-featured sleep-wake and seizure scoring platform to those with access to basic computing resources. This tool will benefit the epilepsy research community as we conduct studies to better characterize and investigate the relationship between sleep, epilepsy, and other comorbidities.

## Data Sharing and Availability

In order to enable the use of the fully-trained models described in this manuscript, all Jupyter Notebooks used for preprocessing and training have been uploaded to GitHub, along with example files for testing importation and viewing file formatting specifications (https://github.com/epilepsylab/SWISC). The full dataset used for the training, validation, and testing splits is available upon reasonable request.

## Funding and Acknowledgements

CURE Epilepsy Award (NPP), NIH R21NS122011 (NPP), NIH K08NS105929 (NPP). We greatly appreciate the input of Matthew Rowan.

