## Supplementary Figure 1; Supplementary Figure 2 for "Automated Classification of Sleep-Wake States and Seizures in Mice"

### Extended Data

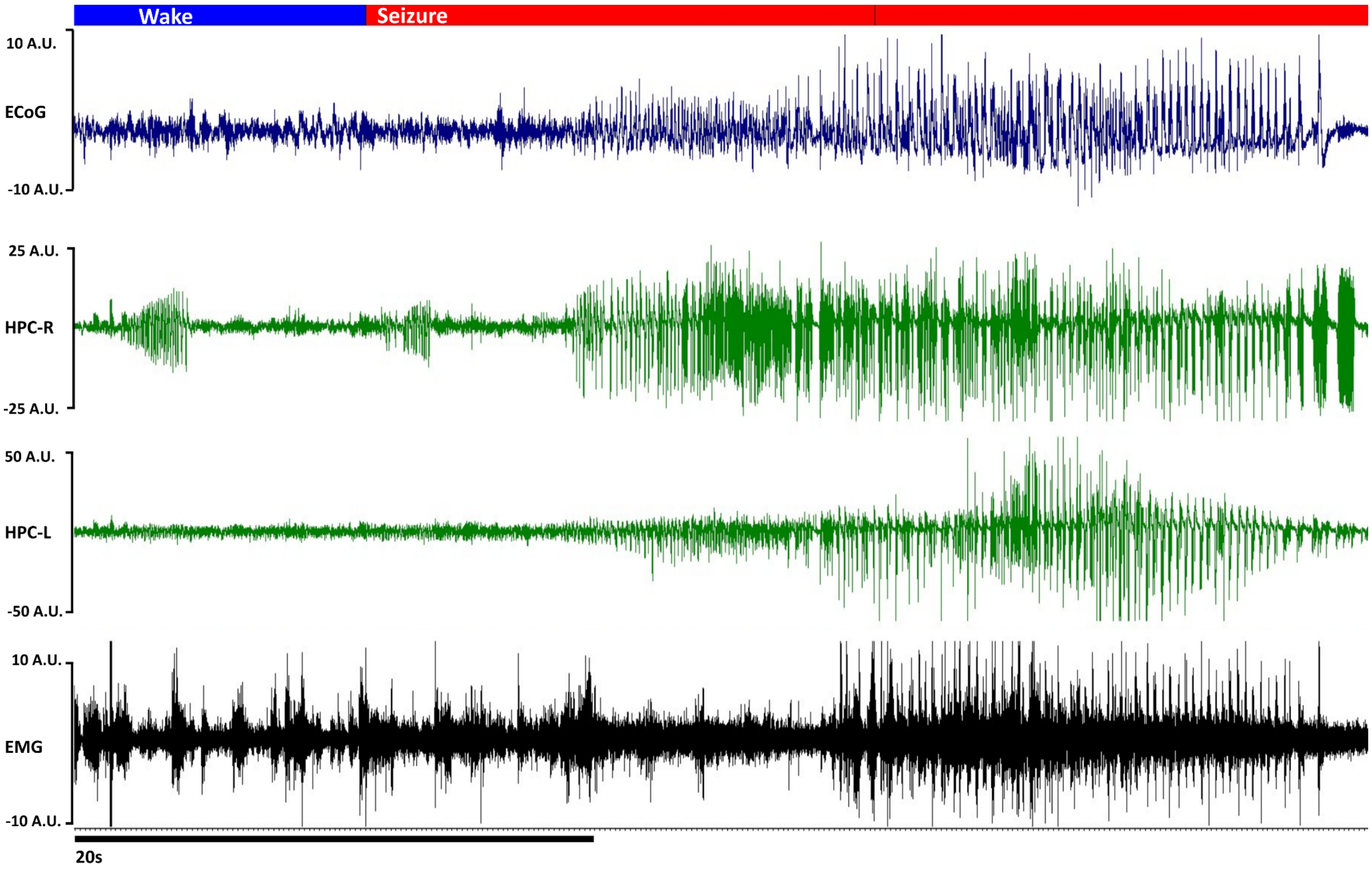


**Supplementary Figure 1.** The seizure from Figure 1.B. is shown magnified for detail and clarity.

| Feature Set | # 1<br>Delta/Theta & RMS EMG<br><br>"DT/RMS" | # 2<br>Statistical Features & RMS EMG<br><br>"Stat/RMS" | # 3<br>FFT Features & RMS EMG<br><br>"FFT/RMS" | # 4<br>Statistical Features, FFT & PSD, & RMS EMG<br><br>"Full" |
|---|---|---|---|---|
| Feature Count | 2 features/channel<br>+ 20 RMS<br>Total: 28 features | 6 features/channel<br>+ 20 RMS<br>Total: 44 features | 7 features/channel<br>+ 20 RMS<br>Total: 48 features | 20 features/channel + 20 RMS<br>Total: 100 features |
| Statistical Measures | - | $\bar{x}$, $\tilde{x}$, σ, v, skewness, kurtosis | - | $\bar{x}$, $\tilde{x}$, σ, v, skewness, kurtosis |
| δ<br>2 - 4 Hz | - | - | Normalized FFT | Normalized FFT & PSD |
| Low-θ<br>4 - 7 Hz | - | - | Normalized FFT | Normalized FFT & PSD |
| High-θ<br>7 - 13 Hz | - | - | Normalized FFT | Normalized FFT & PSD |
| β<br>13 - 30 Hz | - | - | Normalized FFT | Normalized FFT & PSD |
| Low-γ<br>30 - 55 Hz | - | - | Normalized FFT | Normalized FFT & PSD |
| High-γ<br>65 - 100 Hz | - | - | Normalized FFT | Normalized FFT & PSD |
| Ratios | $\frac{\lvert\overline{\text{FFT}(\delta)}\rvert}{\lvert\text{FFT}(\text{Low}-\theta)\rvert}, \frac{\lvert\overline{\text{PSD}(\delta)}\rvert}{\lvert\text{PSD}(\text{Low}-\theta)\rvert}$ | - | $\frac{\lvert\overline{\text{FFT}(\delta)}\rvert}{\lvert\text{FFT}(\text{Low}-\theta)\rvert}$ | $\frac{\lvert\overline{\text{FFT}(\delta)}\rvert}{\lvert\text{FFT}(\text{Low}-\theta)\rvert}, \frac{\lvert\overline{\text{PSD}(\delta)}\rvert}{\lvert\text{PSD}(\text{Low}-\theta)\rvert}$ |
| EMG | 1 sec RMS Binning | 1 sec RMS binning | 1 sec RMS Binning | 1 sec RMS Binning |

**Supplementary Figure 2. Feature Set Selection.**

Testing the feature space dependence of the various classification regimes was accomplished by selecting three groups of features to compare. The first (**DT/RMS**) consisted solely of the four channels' delta/theta ratios in both FFT magnitude and PSD, as well as the full vector of RMS EMG amplitude. This most closely reproduces the features most important to a sleep-wake expert scorer when scoring manually. The second feature set **(Stat/RMS)** included seven statistical features per channel for each epoch, as well as the RMS EMG amplitude. The third feature set (**FFT/RMS)** included only the four channels' Fourier magnitudes in the selected bins (normalized to broadband 2-55 Hz magnitude), the delta/theta ratio, and the RMS EMG. No epoch-level statistical features nor Power Spectral Density were used for Feature Set 2. The final evaluated feature set contained all statistical features, FFT and PSD magnitudes (both normalized to their relative 2 - 55 Hz broadband magnitudes), delta/theta ratios for FFT and PSD, and RMS EMG components of the full 100-feature vector, and is referred to in subsequent figures as the **Full** feature space.
